# ‘Opioid addiction vs. ‘sugar addiction’: Mesocorticolimbic stimulation controls motivation for remifentanil or alternative reward

**DOI:** 10.64898/2026.10.12.758585

**Authors:** Ileana Morales, David Nguyen, Kent C. Berridge

## Abstract

Addiction is characterized by intense incentive motivation that becomes narrowly focused on particular targets. Laboratory studies have modeled intensification, directional targeting, and narrowing of the focus of incentive motivation using paired optogenetic stimulation of central amygdala CeA circuitry, which can create addictive-like motivation focused on either cocaine or on sucrose, as directed by experimenters. However, opioid drugs have been suggested to be different from cocaine in both reward and addictive effects, to be less dependent on mesolimbic incentive salience circuitry, and so may be potentially exempt from such control. To test if opoioid motivation is subject to similar control, CeA ChR2 rats were allowed to freely choose between intravenous remifentanil (2 ug/kg) or sucrose. One group had CeA stimulation selectively paired with earning sucrose, while the other group had stimulation paired with remifentanil. The sucrose-paired CeA ChR2 rats came to nearly exclusively pursue sucrose, ignoring remifentanil. By contrast, remifentanil-paired CeA ChR2 rats specifically pursued remifentanil, ignoring sucrose (control eYFP rats had no overall preference). To confirm whether CeA ChR2 pairing also intensified incentive motivation for remifentanil, as well as controlling its target, a separate CeA ChR2 group chose between two remifentanil options: one paired with CeA Stimulation, and the other unpaired. CeA ChR2 stimulation both focused pursuit specifically on laser-paired remifentanil, and escalated breakpoint for laser-paired remifentanil in a progressive ratio test of motivation intensity. Despite this ability to control motivation for opioid or sucrose, CeA ChR2 laser on its own had only low to moderate incentive value. For example, some CeA ChR2 rats above completely failed to self-stimulate CeA laser by itself. It appears that amygdala-related recruitment of mesocorticolimbic circuitry can transform the incentive salience of opioid drugs, similar to that for cocaine or sucrose, and gate the induction of intense but narrowly focused ‘wanting’ in an addictive-like fashion.

**Significance Statement:** Optogenetic central amygdala (CeA) circuitry in rats can selectively recruit mesolimbic circuitry to arbitrarily direct, amplify, and narrow incentive motivation onto either cocaine or sucrose. However, opioid drugs have additional effects that differ from psychostimulants and may resist such CeA control. We allowed CeA ChR2 rats to rats to freely choose to earn either intravenous remifentanil or sucrose pellet rewards. One group of CeA ChR2 rats had laser paired with sucrose but not remifentanil, whereas another group had laser paired with remifentanil but not sucrose. CeA ChR2 rats that had laser paired with sucrose, exclusively chose sucrose and ignored the opportunity for intravenous remifentanil. By contrast CeA ChR2 rats that had laser paired with remifentanil chose remifentanil and ignored sucrose.

## Introduction

Opioid drugs have rewarding and addictive effects that may be different from psychostimulants like cocaine or methamphetamine. For example, opioid reward and addiction have been suggested to be independent of the mesolimbic dopamine-related circuitry athat mediates incentive salience (1–11). Consistent with that objection, opioid drugs are often taken in different situations from cocaine and amphetamine (1–6, 12).

Opioids are not necessarily more motivationally potent rewards than other alternatives. For example, most rats prefer sucrose or social rewards over an opioid drug in at least some situations, and the presence of alternative rewards can reduce opioid intake (13–16). Lenoir and colleagues found approximately 40% of rats still failed to prefer heroin over saccharin even after extended i.v. heroin experience, and 25% oppositely preferred saccharin over heroin (12). Clinically, many people who use opioids in their teens and 20s may eventually give up drug use by their 40s (17, 18).

Yet opioid addiction, induced in vulnerable individuals, may create more powerful motivation than use by nonaddicted users (12, 19–22). Incentive sensitization has been suggested to cause excessive drug ‘wanting’ in individuals vulnerable to mesocorticolimbic sensitization, and to apply to opioid addiction as well as to other drugs (19, 23). After extended drug experience, rats increase their preference for heroin over saccharin three times more than for cocaine over saccharin (12).

Laboratory studies have modeled the amplification and narrow focusing of incentive salience to produce intense ‘wanting’ similar to incentive sensitization (24–27). Those studies directionally assigned ‘wanting’ using optogenetic channelrhodospsin (ChR2) stimulations of central nucleus of amygdala (CeA) circuitry, which is associatively paired with an experimenter-chosen target such as cocaine, or sucrose, or a shock rod (24–27). CeA pairing was used because amygdala-related circuitry helps assign reward-related and fear-related motivational significance to stimuli in the environment, and CeA is able to direct the targeting of intense incentive motivation (28–39). CeA in particular has striatal-like status in macrosystem frameworks (40–44), and optogenetic or neurochemical stimulations in CeA can generate intense appetitive motivated behaviors (24–27, 32, 33, 45, 46).

Here we assessed whether associative pairing of CeA ChR2 optogenetic stimulation could similarly control the focus and intensity of motivation for an opioid drug, remifentanil. Remifentanil is a fast-acting mu-opioid agonist, approximately 100-times more potent than morphine (47, 48) and is readily self-administered by rats (49, 50). Our results indicate that optogenetic CeA ChR2 pairing can control the induction and directional targeting of addictive-like intense and narrowly focused motivation for remifentanil versus sucrose. These results are consistent with the incentive sensitization theory of addiction, which suggests that addictive-like motivation for drugs, including opioids, can be produced by selective hyper-reactivity of mesocorticolimbic circuitry that generated and assigns excessive incentive salience to particular targets.

## Results

### Experiment 1: Remifentanil vs. Sucrose

CeA ChR2 (**Fig. 1a**) and control CeA eYFP rats chose freely between sucrose pellets or i.v. remifentanil infusions (**Fig. 1b**). For some rats, CeA laser was paired with earning sucrose but not remifentanil, whereas for other rats, CeA laser was paired with remifentanil but not sucrose. CeA ChR2 rats that had laser paired with earning sucrose but not remifentanil came to choose *Laser + Sucrose* almost exclusively while ignoring the opportunity to earn i.v. *Remifentanil Alone* (**Fig. 1b-e**; sucrose + laser group: *F_8,32_*= 4.95, *p* = 0.005, *n* = 5). A 2:1 ratio of *Laser + Sucrose* preference emerged by the second day of 2-choice testing (**Fig S1a**; *t_32_=* 3.12, *p* < 0.03), and continued to grow to a 6:1 preference for *Laser + Sucrose* over *Remifentanil Alon*e by the final RR6 test on day 9 (*t_32_=* 8.33, *p* < 0.0001).

**Figure 1.**
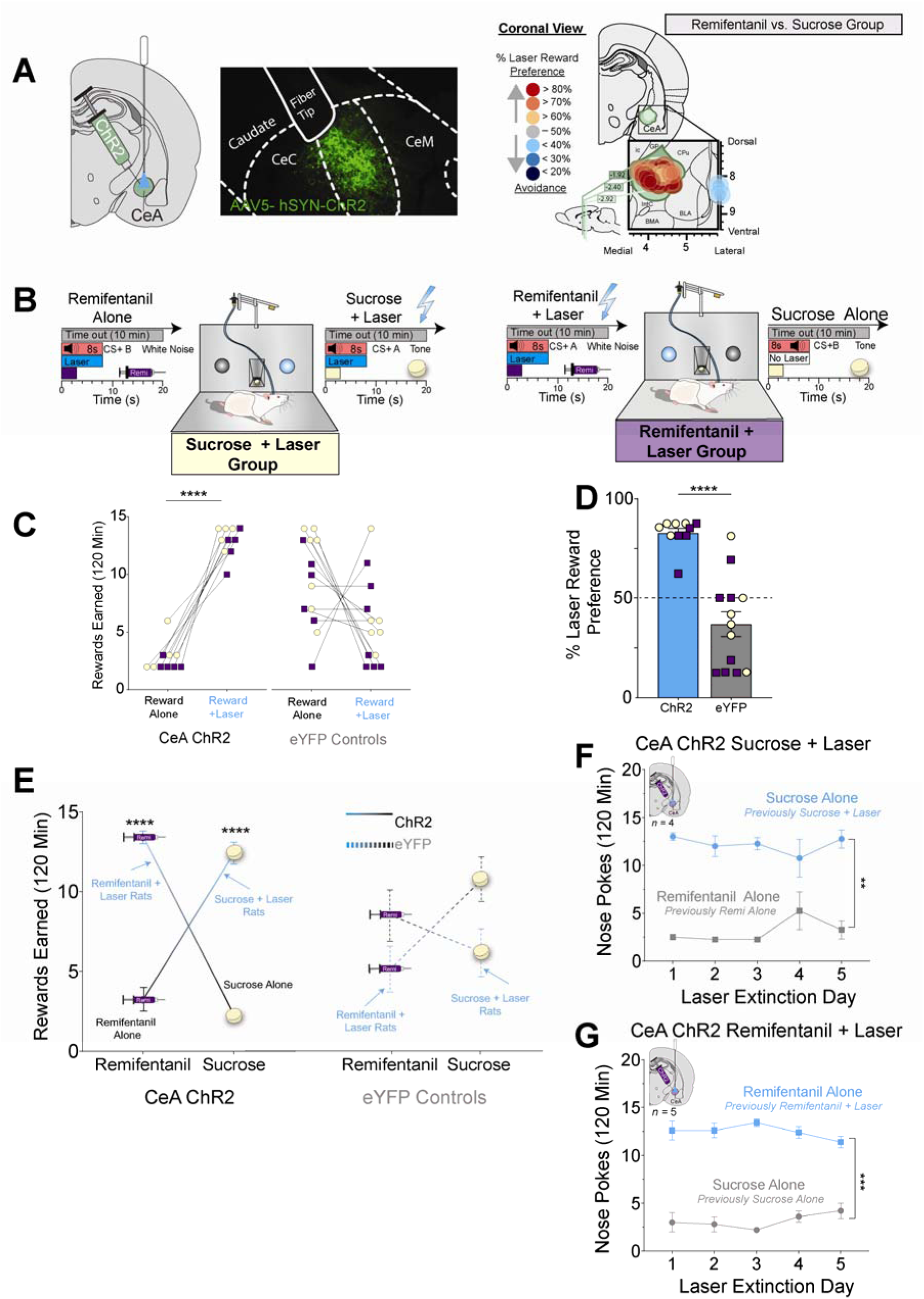
CeA neurons control choice between remifentanil and sucrose. **A)** CeA photomicrograph shows green channelrhodopsin (ChR2; AAV5-hSYN-ChR2-eYFP) expression in CeA. Right: Mapped sites of optic fiber implants for each ChR2 rat in remifentanil vs. sucrose group. Symbol colors in both groups represent the percent preference for an individual rat’s preference for that rat’s laser-paired reward over the non-laser paired reward calculated as the average of the last two sessions. Ic, internal capsule; GP, globus pallidus; CPu, caudate putamen; BLA basolateral amygdala, CeA, central amygdala; IntC, intercalated amygdala; BMA, basomedial amygdala. **B)** Describes remifentanil vs sucrose choice paradigm; s, seconds, min, minutes. **C)** Earned rewards (nose pokes; FR1 schdule) during 120-min remifentanil vs. sucrose choice sessions for individual rats (ChR2: *n* = 10; eYFP: *n* = 13). **D)** Percent preference for laser-paired reward on day 9 (ChR2: *n* = 10; eYFP: *n* = 13). Individuals with laser paired with sucrose in yellow circles, and remifentanil in purple squares. **E)** Total number of rewards sucrose and remifentanil rewards earned on day, separated by reward type, and virus group. ChR2 rats in solid lines; eYFP rats in dotted lines. ChR2: remifentanil + laser *n = 5*, sucrose + laser *n = 5*; eYFP: remifentanil + laser *n = 7*; sucrose + laser *n* = 6 **F, G)** Reward preference between remifentanil and sucrose persists after CeA neuron excitation pairing is placed in extinction. Data presented as mean ± SEM; \*\**p* < 0.01, \*\*\**p* < 0.001, \*\*\*\**p* < 0.0001

Conversely, the other group of CeA ChR2 rats that had laser paired with remifentanil but not sucrose chose *Laser + Remifentanil* primarily while ignoring the opportunity to earn *Sucrose Alone* (**Fig. 1c-e**; remifentanil + laser group: *F*(*_1,4_*) = 50.99, *p* = 0.002, *n* = 5). Their preference for *Laser + Remifentanil* over *Sucrose Alon*e grew to a 4:1 ratio by day 9 (**Fig. S1b**).

By contrast, control eYFP rats with optically inactive virus in CeA (eYFP) rats chose equally between sucrose and remifentanil as a group, regardless of which reward was laser paired, and this differed from ChR2 rats (**Fig. 1c-e, S1c-d**; eYFP remifentanil + laser group: *F_8,48_*= 1.51, *p* = 0.18, *n* = 7; eYFP sucrose + laser group: *F_8,40_*= 1.03, *p* = 0.18, *n* = 6). For both groups, no sex differences were detected in the magnitude of laser-induced preference (**Fig. S1e**; female 4:1 ratio preference; males 5:1 ratio; *F_8,64_*= 1.00, *p* = 0.44, *n* = 6 females, *n =*4 males).

#### Cea ChR2-induced preferences persist after laser is discontinued

To assess whether a laser-established preference would persist if CeA laser pairing were discontinued, an additional 5 days of laser-extinction tests were run after laser-accompanied 2-choice tests had finished. Despite the discontinuation of laser stimulation, previous laser-established preferences persisted nearly unchanged for the next 5 no-laser days. The group of CeA ChR2 rats that previously had *Laser + Sucrose* preference over *Remifentanil Alone* persisted in choosing the now *Sucrose Alone* option over *Remifentanil Alone* by a 4:1 ratio through day 5 (**Fig. 1f**; laser main effect; *F_1,3_* = 34.41, *p* < 0.001; *n* = 4). Similarly, CeA ChR2 rats CeA ChR2 rats with a previous *Laser + Remifentanil* preference over *Sucrose Alon*e, still continued to choose the now *Remifentanil Alon*e option over *Sucrose Alon*e by a 3:1 ratio (**Fig. 1g**; laser main effect; *F_1,4_* = 83.65, *p* < 0.001; *n* = 5).

### Is laser the goal? Laser self-stimulation in CeA ChR2 rats

#### Rod/Spout-touch laser self-stimulation

In the rod-touch or spout-touch task, rats were classified as high self-stimulators if they earned >50 laser illuminations per session, and made at least 2x as many contacts on the laser-delivering active rod or spout as on the inactive control spout (473 nm; 1-2 mW, 25 Hz; 1-s or 8-s bin duration depending on day). Rats were classified as moderate self-stimulators if they earned 10-49 laser illuminations per session, and made at least 2x as many contacts on the active rod/spout as on the inactive control spout. Rats were classified as failures to self-stimulate if they earned <10 illuminations or failed to touch their laser spout twice as often as the control spout. All rats were classified on Day 1 and re-tested for reliability of self-stimulation on Days 2 and 3, both before and after 2-choice tasks.

##### 1- sec laser bins: more reliable CeA ChR2 self-stimulation

###### Experiment 1: Remifentanil vs. Sucrose Group

*1-s bin self-stimulation*. CeA ChR2 rats reliably self-stimulated for 1-s laser bins when tested before (**Fig. 2a**; *F*_1,18_ = 8.47, *p = 0.009*, *n* = 8 CeA ChR2 rats; *Control Spout contacts:* 8.3 ± 3.5, *Laser Spout:* 45.7 ± 17.7) and again after Remi vs. Sucrose 2-Choice tests (*F*_1,18_ = 15.67, *p < 0.001*, *n* = 8 CeA ChR2 rats; *Control Spout contacts:* 13.0 ± 2.6, *Laser Spout:* 506.3 ± 158.1), consistent with several previous reports (25, 26, 45, 46, 51–55). There was still individual heterogeneity within those groups, as 12.5% (before) and 87.5% (after) of CeA ChR2 rats self-stimulated at high levels, 50% (before) and 0% (after) of CeA ChR2 rats self-stimulated at moderate level, and 50% (before) and 12.5% of CeA ChR2 (after) failed to self-stimulate, again similar to previous reports (25, 26, 45, 46).

**Figure 2.**
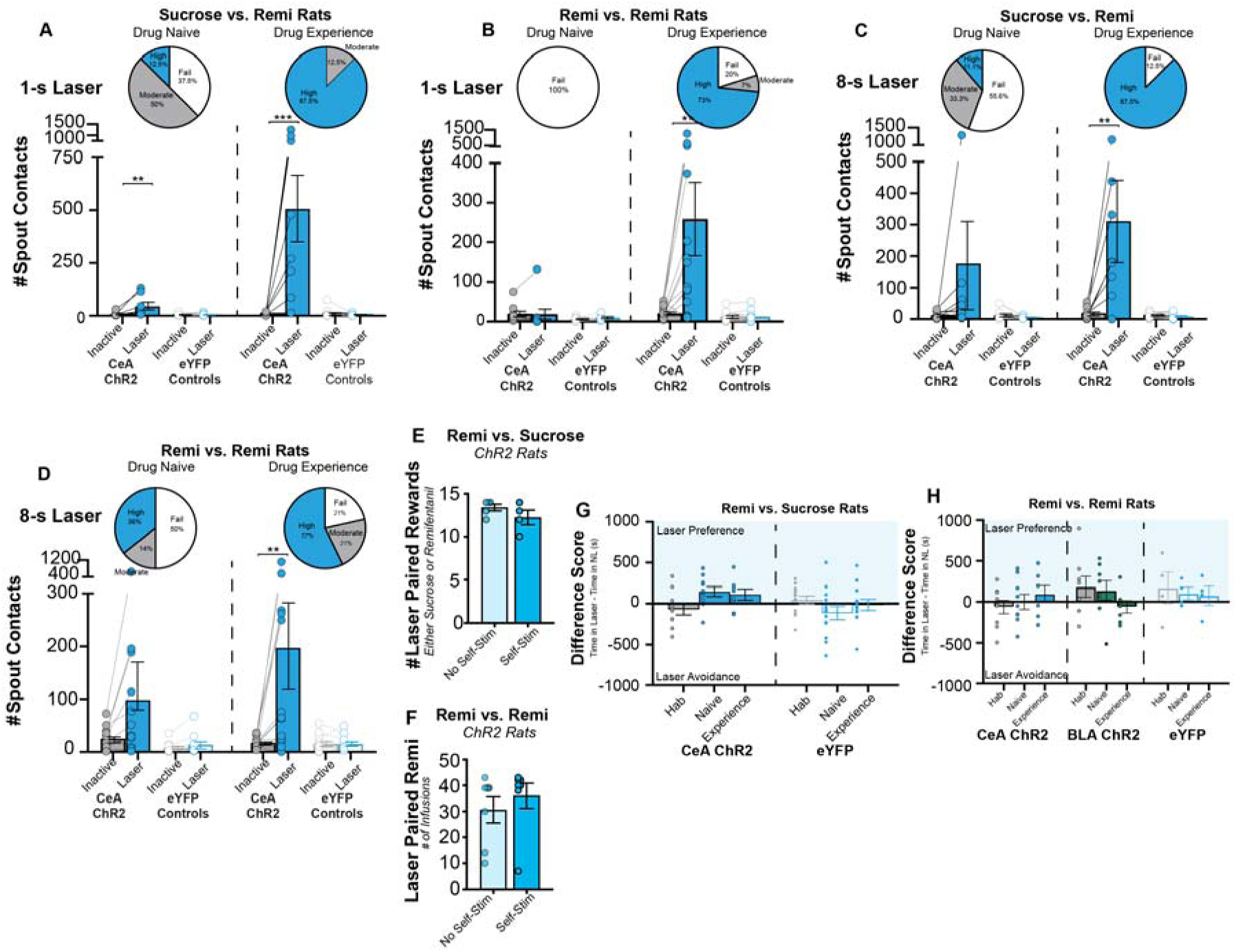
CeA neurons do not reliably promote self-stimulation. **A-D)** Active spout-based laser self-administration task where touching a laser spout earned laser stimulation (25 Hz, 8-s or 1-s duration, 1-2 mW) and touching a separate inactive spout earned nothing. **A)** 1-s laser stimulations produce reliable self-stimulation in CeA ChR2 rats from remifentanil vs. sucrose experiments when drug naïve (left; ChR2: *n*=8, eYFP: *n* = 12) and after drug experience (right; ChR2: *n*=10, eYFP: *n* = 12). **B)** No reliable self-stimulation in CeA ChR2 rats from remifentanil vs. remifentanil experiments when drug naïve (left; ChR2: *n*=10, eYFP: *n* = 9). When tested again after drug experience (right; ChR2: *n*=14, eYFP: *n* = 13), CeA ChR2 rats significantly self-stimulated for 1-s laser. **C)** Total laser self-administrations earned by rats from remifentanil vs. sucrose experiments when drug naïve (left; ChR2: *n*=9, eYFP: *n* = 11) and after drug experience (right; ChR2: *n*=8, eYFP: *n* = 10). CeA ChR2 rats do not reliably self-stimulate for 8-s when drug naïve, but self-stimulate after remifentanil/sucrose experience. **D)** Total laser self-administrations earned by rats from remifentanil vs. remifentanil 2-choice experiments when drug naïve (left; ChR2: *n*=14, eYFP: *n* = 10) and after drug experience (right; ChR2: *n*=14, eYFP: *n* = 13). CeA ChR2 rats again do not reliably self-stimulate for 8-s when drug naïve, but self-stimulate after remifentanil/sucrose experience. **E)** Self-stimulating CeA ChR2 rats earned equal laser-paired rewards compared to non-stimulating CeA ChR2 rats in remi vs. sucrose experiments (non-self-stim: *n* = 5; self-stim: *n* = 4). **F)** Self-stimulating CeA ChR2 rats earned equal laser-paired remifentanil compared to non-stimulating CeA ChR2 rats in remi vs. remi experiments (non-self-stim: *n* = 7; self-stim: *n* = 7). **G, H)** No place-based self-stimulation in any rats from Experiment 1 (ChR2: *n* = 9; eYFP: *n* = 13) or Experiment 2 (CeA ChR2: *n* = 9; BLA ChR2: *n* = 4 ; eYFP: *n* = 8). Data presented as mean ± SEM; \*\**p* < 0.01, \*\*\**p* < 0.001.

Control eYFP rats failed as a group to self-stimulate in the spout task, made equal contacts to the laser spout and control spout (**Fig 2a**; Before remi vs. sucrose: Control spout: 7.2 ± 1.9, Laser Spout: 6.5 ± 1.6, *t_18_ =* 0.08, *p* = 0.99, *n* = 12; After remi vs. sucrose: Control spout: 13.0 ± 6.1, Laser Spout: 5.6 ± 1.8, *t_18_ =* 0.09, *p* = 0.99, *n* = 12) and differed from CeA ChR2 rats a both times tested (Before remi vs. sucrose: *F*_1,18_ = 6.01, *p = 0.025;* After remi vs. sucrose: *F*_1,18_ = 15.15, *p = 0.001)*.

###### Experiment 2: Remifentanil 2-Choice Rats

*Initial 1-s bin Self-Stimulation*. In contrast to CeA ChR2 rats from Experiment 1, CeA ChR2 rats that were tested in the Remifentanil 2-Choice task (Experiment 2) as a whole group failed to significantly self-stimulate laser (**Fig. 2b**; *F*_1,18_ = 0.31, *p = 0.59*, *n* = 10 CeA ChR2 rats; *Control Spout contacts:* 18.8 ± 6.8, *Laser Spout:* 18.0 ± 12.7). 100% of CeA ChR2 rats failed to meet criteria for self-stimulation. Control eYFP rats also failed as a group to self-stimulate in the spout task, made equal contacts to the laser spout (9.1± 2.8) and control spout (5.4± 1.9) and did not differ from CeA ChR2 rats (*F*_1,17_ = 1.17, *p = 0.29*, *n* = 9 eYFP rats).

*Post Opioid/Sucrose 1-s bin Self-Stimulation*. After the end of 2-choice tasks that associatively paired CeA laser with either remifentanil or sucrose, rats were retested again for self-stimulation performance. After experience with remifentanil paired CeA laser, the CeA ChR2 group as a whole now did robustly self-stimulate 1-s laser bins (**Fig. 2b** *; F*_1,25_ = 6.24, *p = 0.02*, *n* = 15 CeA ChR2 rats; *laser spout contacts:* 258.6 ± 91.6; *control spout contacts*: 20.1 ± 3.7; *t_25_* = 3.6*, p =* 0.003). Individually, 11 of 15 CeA ChR2 rats (73%) now met criteria for high self-stimulation, 1 rat (7%) was a moderate self-stimulator), and only 3 rats failed to self-stimulate (20%).

By comparison, control eYFP rats still failed to still self-stimulate both as a group and as individuals (**Fig. 2b**; laser spout contacts: 11.5 ± 3.9; control spout contacts: 11.6 ± 3.7; *t_25_* = 0.000000*, p =* 0.99), and thus the CeA eYFP control group now differed from the CeA ChR2 group in self-stimulation (*F*_1,25_ = 6.83, *p =* 0.02, *n* = 13 eYFP). We conclude that 1-s laser durations more reliably produce laser self-stimulation, although some heterogeneity is observed across groups.

##### 8-sec Laser: Little CeA ChR2 self-stimulation initially, more after reward association

###### Experiment 1: Remifentanil vs. Sucrose Group

*Initial 8-s bin Self-Stimulation*. When tested with 8-s laser bins before Remi vs. Sucrose2-choice tests, the Experiment 1 CeA ChR2 group as a whole failed to significantly self-stimulate laser (**Fig. 2c**; *F*_1,18_ = 1.77, *p = 0.20*, *n* = 9 CeA ChR2 rats; *Control Spout contacts:* 12.1 ± 3.7, *Laser Spout:* 176.4 ± 139.3). However, 1 CeA ChR2 rat met criteria for self-stimulation, 3 CeA ChR2 rats (33.3%) were moderate self-stimulators, and 5 CeA ChR2 rats (55.6%) of rats completely failed to self-stimulate in 8-sec bins.

Control eYFP rats also failed as a group to self-stimulate in the spout task, made equal contacts to the laser spout (5.8± 1.0) and control spout (12.4± 4.4) and did not differ from CeA ChR2 rats (**Fig. 2c**; *F*_1,18_ = 1.83, *p = 0.19*, *n* = 9 ChR2, *n* = 11 eYFP rats). We conclude that 8-sec CeA laser stimulation is relatively inconsistent as an intrinsic incentive, given that a greater proportion of CeA ChR2 rats initially failed completely to self-stimulate than met high self-stimulation criteria (and a comparably-sized intermediate group met only low self-stimulation criterion.

*Post Opioid/Sucrose 8-s bin Self-Stimulation*. After the end of 2-choice tasks that associatively paired CeA laser with either remifentanil or sucrose, rats were retested again for self-stimulation performance. After experience with remifentanil/sucrose-paired CeA laser, the CeA ChR2 group as a whole now did robustly self-stimulate 8-s laser bins (**Fig. 2c***; F*_1,18_ = 8.05, *p = 0.01*, *n* = 8 CeA ChR2 rats; *laser spout contacts:* 312.4 ± 130.1; *control spout contacts*: 19.4 ± 6.5; *t_18_* = 3.62*, p =* 0.004). Individually, 7 of 8 CeA ChR2 rats (87%) now met criteria for high self-stimulation, none were moderate self-stimulators, and only1 CeA ChR2 rat still failed to self-stimulate (12.5 %).

By comparison, control eYFP rats still failed to still self-stimulate both as a group and as individuals (**Fig. 2c**; laser spout contacts: 9.8 ± 2.8; control spout contacts: 13.5 ± 2.6; *t_18_* = 0.05*, p =* 0.99), and thus the CeA eYFP control group now differed from the CeA ChR2 group in self-stimulation (*F*_1,18_ = 8.55, *p =* 0.009, *n* = 8 ChR2, *n* = 12 eYFP). We conclude that prior association with either remifentanil or sucrose markedly enhances the incentive value of 8-sec CeA laser bin stimulation for most CeA ChR2 rats, adding significantly its initial inherent incentive value.

###### Experiment 2: Remifentanil 2-Choice Rats

*Initial 8-s bin Self-Stimulation*. When tested before 2-choice tests, this separate group of CeA ChR2 rats (n = 14 total) as a whole failed to significantly self-stimulate 8-s CeA laser bins (**Fig. 2d**; *F*_1,22_ = 1.90, *p = 0.18*), although as a group they made 4x as many touches on the laser-paired spout (98.6 ± 45.5) as on the control spout (25.1 ± 5.6). Individually, 5 of 14 CeA ChR2 rats (36%) met criteria for high self-stimulation, 2 CeA ChR2 rats (14%) were low self-stimulators, and 7 (50%) completely failed to self-stimulate 8-s CeA laser.

Control eYFP rats also failed to self-stimulate 8-s laser, making equally low numbers of touches on the laser spout (13.6 ± 6.6) and control spout (8.0 ± 3.3). Overall, control eYFP rats as a group did not differ from the group of CeA ChR2 rats in initial levels of 8-sec laser self-stimulation (**Fig. 2d**; *F*_1,22_ = 2.88, *p = 0.10*, *n* = 10 eYFP). We conclude again that 8-s CeA ChR2 laser stimulation has relatively weak or inconsistent intrinsic incentive value.

*Post Opioid 8-s bin Self-Stimulation*. Retested after 2-choice tests where 8-s laser became associated with remifentanil infusions from one instrumental option, drug experienced CeA ChR2 rats as a group now did significantly self-stimulate 8-s CeA laser illuminations (**Fig. 2d***; F*_1,25_ = 4.5, *p = 0.04*, *n* = 14 ChR2). On average, CeA ChR2 rats now made 12x as many contacts to the laser paired spout (197.4 ± 82.3) relative to the control spout (17.0 ± 2.9; *t_25_* = 3.08*, p =* 0.01). Individually, 8 of 14 CeA ChR2 rats (57%) now self-stimulated at high levels, 3 CeA ChR2 rats (21%) met criteria for low self-stimulation, and only 3 CeA ChR2 rats (21%) still failed to self-stimulate.

By comparison, control eYFP rats failed to self-stimulate as a group. Individually, only 1 eYFP rat met criteria for moderate self-stimulation, and 12 eYFP rats completely failed to self-stimulate, making equal contacts between the control spout (14.7 ± 5.0) and laser spout (15.1 ± 4.3). The control eYFP group thus differed from the CeA ChR2 group in laser self-stimulation (**Fig. 2d**; *F*_1,25_ = 4.5, *p =* 0.04, *n* = 13 eYFP). We again conclude that a prior association of CeA laser with remifentanil can increase the self-stimulation incentive value of CeA laser by itself above its initial incentive value for most CeA ChR2 rats.

##### Experiment 1 &2: Effect of Initial self-stimulation status on remifentanil or sucrose pursuit

The analyses above indicate that 8-sec CeA ChR2 stimulation initially has relatively little incentive value for most rats, as measured by 8-s laser self-stimulation, but gains increased value after association with remifentanil or sucrose rewards. However, a minority 22% - 36% of CeA ChR2 individuals did self-stimulate 8-s laser at high levels even initially, and an additional 14% - 36% initially self-stimulated at least at moderate levels. Thus approximately one-half of our CeA ChR2 groups showed some level of initial laser self-stimulation, while the remainder did not.

This raises the question of how much attempts to self-stimulate laser contributed to specific pursuit of laser-paired remifentanil or laser-paired sucrose in subsequent 2-choice or breakpoint tasks in Experiments 1 and 2.

*Experiment 1: Remifentanil vs. Sucrose*. To answer the question of whether CeA ChR2 rats pursued a laser-paired reward over a non-paired reward as an attempt to gain laser self-stimulation, we directly compared individual preference ratios for pursuit of their laser-paired sucrose or laser-paired remifentanil over the non-paired alternative reward in CeA ChR2 individuals that initially self-stimulated 8-s CeA laser versus in CeA ChR2 individuals that completely failed to initially self-stimulate 8-s CeA laser. Results indicated that initial laser self-stimulation propensity did not measurably contribute to CeA ChR2-induced pursuit of laser laser-paired sucrose or laser-paired remifentanil. That is, initially self-stimulating CeA ChR2 rats did not earn more laser-paired sucrose or laser-paired remifentanil than other CeA ChR2 rats that failed to initially self-stimulate (**Fig 2e**; Self-stimulators: 12.3 ± 0.9, *n* = 4; Failures to self-stimulate:13.4 ± 0.4; *n* = 5; *t_7_* = 1.31*, p =* 0.23). Consequently, there was no significant correlation between initial laser self-stimulation score and preference ratio for laser-paired sucrose or laser-paired remifentanil in CeA ChR2 rats (Pearson’s correlation, r_9_ = 0.09, p = 0.81).

*Experiment 2: Remifentanil vs. Remifentanil*. We similarly compared individual preference ratios for pursuit of their laser-paired remifentanil over non-paired remifentanil in CeA ChR2 individuals that initially self-stimulated 8-s CeA laser versus in CeA ChR2 individuals that initially failed to self-stimulate. Again, initial self-stimulation propensity did not detectably contribute to intensifying and narrowing motivation toward the Laser + Remifentanil option. CeA ChR2 rats that initially failed to self-stimulate CeA laser showed virtually as strong pursuit of their laser-paired remifentanil option as other CeA ChR2 rats that did self-simulate made equal laser-paired reward responses, with no statistical difference between the groups (**Fig 2f**; Self-stimulators: 36 ± 4.9, *n* = 7; Failures to self-stimulate: 30.6 ± 5.0, *n* = 7; *t_12_* = 0.77*, p =* 0.45). Again, therefore, there was no detectable correlation between laser self-stimulation score and preference for laser-paired remifentanil in CeA ChR2 rats (Pearson’s correlation, r_14_ = 0.15, *p* = 0.61).

##### Unreliable Place-Based Self-Stimulation

*Experiment 1: Remifentanil vs. Sucrose Rats*. In the place-based task, rats could earn laser illuminations by entering a designated chamber, and laser continued to cycle as long as the rats remained in that chamber (25 Hz; 3-sec ON, 8 sec OFF cycle). CeA ChR2 rats that were tested in the remifentanil vs. sucrose task self-stimulated in the place-based task at moderate levels (**Fig. 2g**; CeA ChR2 Group: Habituation Score: −69.0 ± 70.8; Drug Naïve Score: 144.7 ± 60.3; Drug Experience Score: 108.5 ± 67.0, n = 10; *F_2,40_ =* 5.36, *p* = 0.01). However, when individual test days were compared, no single day was statistically significant (Habituation vs. naïve: *t_40_= 2.49, p = 0.052,* Habituation vs experience: *t_40_= 1.75, p = 0.26,* Naïve vs. Experience: *t_40_= 0.56, p = 0.99)*.

Similarly, eYFP control rats failed to self-stimulate at any point tested (**Fig. 2g**; CeA eYFP Group: Habituation Score: 42.2 ± 48.16; Drug Naïve Score: −115.4 ± 83.7; Drug Experience Score: −16.5 ± 67.4; Habituation vs. naïve: *t_40_= 2.49, p = 0.052,* Habituation vs experience: *t_40_= 1.75, p = 0.26,*Naïve vs. Experience: *t_40_= 0.56, p = 0.99,* n = 13).

*Experiment 2: Remifentanil 2-choice rats*. Similar to rats from Experiment 1: CeA ChR2 rats that went through Remifentanil + Laser vs. Remifentanil Alone 2-Choice testing failed to self-stimulate in the place-based task, spending equal time in the no-laser chamber, and laser-delivering chamber, both before and after experience with i.v. remifentanil (**Fig. 2h**; CeA ChR2 Group: Habituation Score: −87.3 ± 87.8; Drug Naïve Score: 27.8 ± 99.7; Drug Experience Score: 160.2 ± 96.4, n = 9; *F_4,33_ =* 1.18, *p* = 0.34).

Similarly, BLA ChR2 rats and control eYFP virus rats failed to show either place-based self-stimulation or avoidance in the place-based task (**Fig. 2h** BLA ChR2 Group: Habituation Score: 170.3 ± 193.6, Drug Naïve Score: 103.3 ± 77.3, Drug Experience Score: 76.5 ± 116.8, n = 4; eYFP Control Group: Habituation Score: −63.4 ± 81.0, Drug Naïve Score: 1.03 ± 91.95, Drug Experience Score: 97.0 ± 103.1, n =8; experience x virus site interaction: *F_4,33_ =* 1.18, *p* = 0.34). A potential reason why the results of the place-based task might differ from those of the spout self-stimulation task is the fact that the cumulative duration of laser illumination (seconds of total laser per minute) was ∼150% greater than in the spout-based task. It is possible that longer laser durations in the place-based task exceeded an optical level for self-stimulation.

### Altered ‘wanting’ but not ‘liking’: CeA ChR2 stimulation does not enhance sucrose hedonic impact

We tested whether CeA ChR2 stimulation that increased ‘wanting’ for laser-paired sucrose also increased the hedonic impact of ‘liking’ of sucrose taste in CeA ChR2 rats that previously pursued only Laser + Sucrose in 2-choice tests (**Fig. 3a & 3b**). In the absence of laser stimulation, intraoral sucrose infusions elicited primarily positive ‘liking’ reactions (i.e., rhythmic midline tongue protrusions, lateral tongue protrusions, and paw licks) in both CeA ChR2 rats and control eYFP rats (**Fig. 3c**; ChR2 ‘liking’ Laser OFF: 14.9 ± 1.9; eYFP ‘liking’ Laser OFF: 12.0 ± 2.0), and few or no aversive ‘disgust’ reactions (ChR2 ‘disgust’ Laser OFF: 1.3 ± 0.5; eYFP ‘disgust’ Laser OFF: 3.1 ± 1.0; CeA ChR2: *F_1,4_ =* 51.36, *p* = 0.002, CI = 8.4, 19.1, *n =* 5; CeA eYFP: *F_1,5_ =* 12.58, *p* = 0.02, CI = 2.7, 17.0, *n =* 6). Laser CeA stimulation in CeA ChR2 rats failed to increase the number of positive ‘liking’ reactions elicited by sucrose, despite having previously increased ‘wanting’ for sucrose in the same rats (**Fig 3c & 3d**; Positive hedonic reactions CeA ChR2 Laser On: total number 15.5 ± 1.8; rhythmic midline tongue protrusions: Laser-OFF: 6.2 ± 2.0, Laser-ON: 6.7 ± 2.3; *t*_4_ = 0.77, *p* = 0.49; paw licks: Laser-OFF: 5.1 ± 1.6, Laser-ON: 4.7 ± 1.6; *t*_4_ = 0.39, *p* = 0.71; lateral tongue protrusions: Laser-OFF: 3.6 ± 0.9, Laser-ON: 4.1 ± 0.5; *t*_4_ = 0.57, *p* = 0.60). The few negative ‘disgust’ reactions to sucrose were also unchanged by laser stimulation in CeA ChR2 rats (ChR2 ‘disgust’ Laser On: 1.6 ± 0.8; *F_1,4_ =* 0.29, *p* = 0.62).

**Figure 3.**
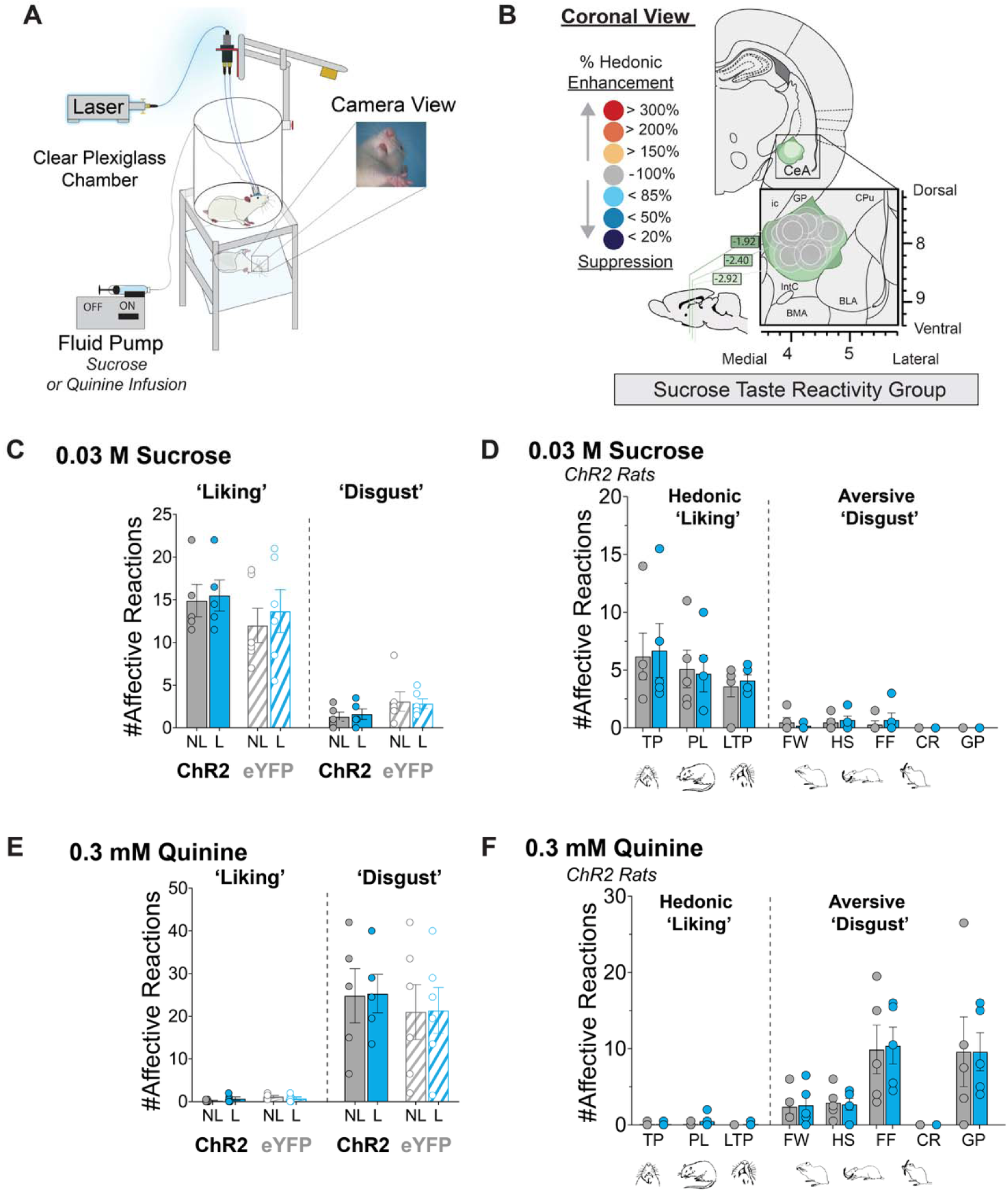
CeA neuron stimulation does not alter hedonic impact for sweet or bitter reward. Depicts taste reactivity test. Rats with stimulus delivery tube (PE-10) tubing connected to oral cannula are placed in Plexiglass chamber. An angled mirror below the clear floor of chamber tilts the ventral view of rat’s face to a video camera recording orofacial expressions. Sucrose (0.03M) or quinine (0.3 mM) infusions (1 mL) are infused directly into rat’s mouth via intraoral cannula at a rate of 1 mL/min via microinfusion pump. **B)** Coronal map depicts anatomical placements and behavioral effects of CeA ChR2 optic fibers in CeA (circles). Color of symbols represents functional hedonic effect on taste reactivity to sucrose of laser illumination at each side: within subjects % increase in sucrose-elicited ‘liking’ reactions by laser illumination over baseline measured to sucrose infusion without laser. **C)** Sucrose CeA ChR2 (solid bars; *n* = 5) and eYFP (striped bars; *n* = 6) affective reactions. No change in total number of positive ‘liking’ reactions (left) and aversive ‘disgust’ expression during no laser conditions (grey bars) and laser stimulation (blue bars). **D)** Separate component ‘liking’ and ‘disgust’ reactions to intraoral infusions of 0.03M sucrose are unchanged in CeA ChR2 rats (*n* = 5) at baseline (grey bars) vs. laser stimulation (blue bars). **E)** No change in total number of affective reactions to quinine of CeA ChR2 rats (solid bars; *n* = 5) and eYFP controls (striped bars; *n* = 6). Blue shows reactions during laser on vs. no laser (grey bars) conditions of the same rats. **F)** Separate component reactions to quinine within hedonic ‘liking’ and aversive ‘disgust’ categories emitted by CeA ChR2 rats (*n* = 5) in laser on (blue) vs. no laser (grey) conditions. Separate component reactions in positive hedonic category are unpacked as: tongue protrusions, TP; paw licks, PL; lateral tongue protrusions, LTP. Separate component reactions in negative aversive category are unpacked as: facewashes, FW; head shakes, HS; forelimb flails, FF; chin rubs, CR; gapes, GP.). Data presented as mean ± SEM.

In absence of laser stimulation, bitter quinine infusions elicited primarily negative ‘disgust’ reactions from both CeA ChR2 rats and eYFP control rats (i.e., gapes, forelimb flails, headshakes, and chin rubs) (**Fig 3e**; ChR2 ‘disgust’’ Laser OFF: 28.4 ± 6.4; eYFP ‘disgust’ Laser OFF: 21.0 ± 6.4; CeA ChR2: *F_1,4_ =* 20.91, *p* = 0.01, CI = −39.5, −9.6, *n =* 5; CeA eYFP: *F_1,5_ =* 11.92, *p* = 0.02, CI = −32.3, −5.2, *n =* 6), and nearly zero positive ‘liking’ reactions (ChR2 ‘liking’ Laser OFF: 0.2 ± 0.1; eYFP ‘liking’ Laser OFF: 1.2 ± 0.2).Laser CeA stimulation in CeA ChR2 rats failed to reduce the number of ‘disgust’ reactions to quinine (**Fig 3e**; CeA ChR2: ‘disgust’ Laser ON: 25.3 ± 4.5), and failed to increase the low number of positive ‘liking’ reactions to quinine in CeA ChR2 rats (Total number of positive hedonic reactions: Laser ON: 0.7 ± 0.4; *F_1,4_ =* 0, *p* = 0.99).

Similarly, in eYFP control rats CeA laser illumination failed to alter ‘liking’ or ‘disgust’ reactions to either sucrose (**Fig 3c**; eYFP ‘liking’ Laser On: 13.7 ± 20.5; eYFP ‘disgust’ Laser On: 2.8 ± 0.6*; F_1,5_ =* 5.2, *p* = 0.08) or quinine (CeA eYFP ‘disgust’ Laser ON: 21.3 ± 5.4, ‘liking’ Laser ON: 0.8 ± 0.3, *F_1,5_ =* 0.06, *p* = 0.81). Positive ‘liking’ reactions of eYFP controls remained similar to CeA ChR2 rats for both tastes (*F_1,9_ =* 0.66, *p* = 0.44), as did the number of aversive ‘disgust’ reactions to quinine (*F_1,9_ =* 0.001, *p* = 0.97). Overall, these results indicate that CeA laser stimulation in CeA ChR2 rats fails to enhance ‘liking’ reactions, despite its powerful control of ‘wanting’ for laser-paired sucrose.

### Experiment 2: Laser + Remifentanil vs. Remifentanil Alone Choice Task

A separate group of CeA ChR2 rats chose between two i.v. remifentanil options, with only one option accompanied by CeA ChR2 laser photoexcitation and the other option delivered without laser (**Fig. 4a**; 1-3 mW; 25 Hz, 8s). CeA ChR2 rats overwhelmingly chose *Laser + Remifentanil,* while relatively ignoring *Remifentanil Alone* (**Fig. S2a**.; *F*_9,162_ = 7.97, *p* < 0.0001, *n =* 19), reaching a 2:1 ratio preference for *Laser + Remifentanil* over *Remifentanil Alone* by Day 3 of FR1 testing (*t*_162_ = 3.50, *p* = 0.006, *d =* 0.92, CI = 1.9 to 18.0), and a 4:1 preference by Day 10 (*t*_162_ = 8.83, *p* < 0.0001, *d =* 2.04, CI = 17.0, 33.1). Subsequently, as effort requirements rose over Days 11 to 15 (FR1, FR2, FR4, RR4, RR6), CeA ChR2 rats maintained a 4:1 *Laser + Remifentanil* preference (**Fig 4b-c**.; *F*_9,139_ = 17.69, *p* < 0.0001, *n =* 18) by day 11 (FR2 ratio; *t*_156_ = 4.10, *p* < 0.0007, *d* = 2.1, CI = 14.7, 82.1), remaining at a 4:1 preference on the final RR6 ratio on Day 15 (*t*_156_ = 7.73, *d* = 1.96, CI = 59.3, 128.6, *p* < 0.0001).

**Figure 4.**
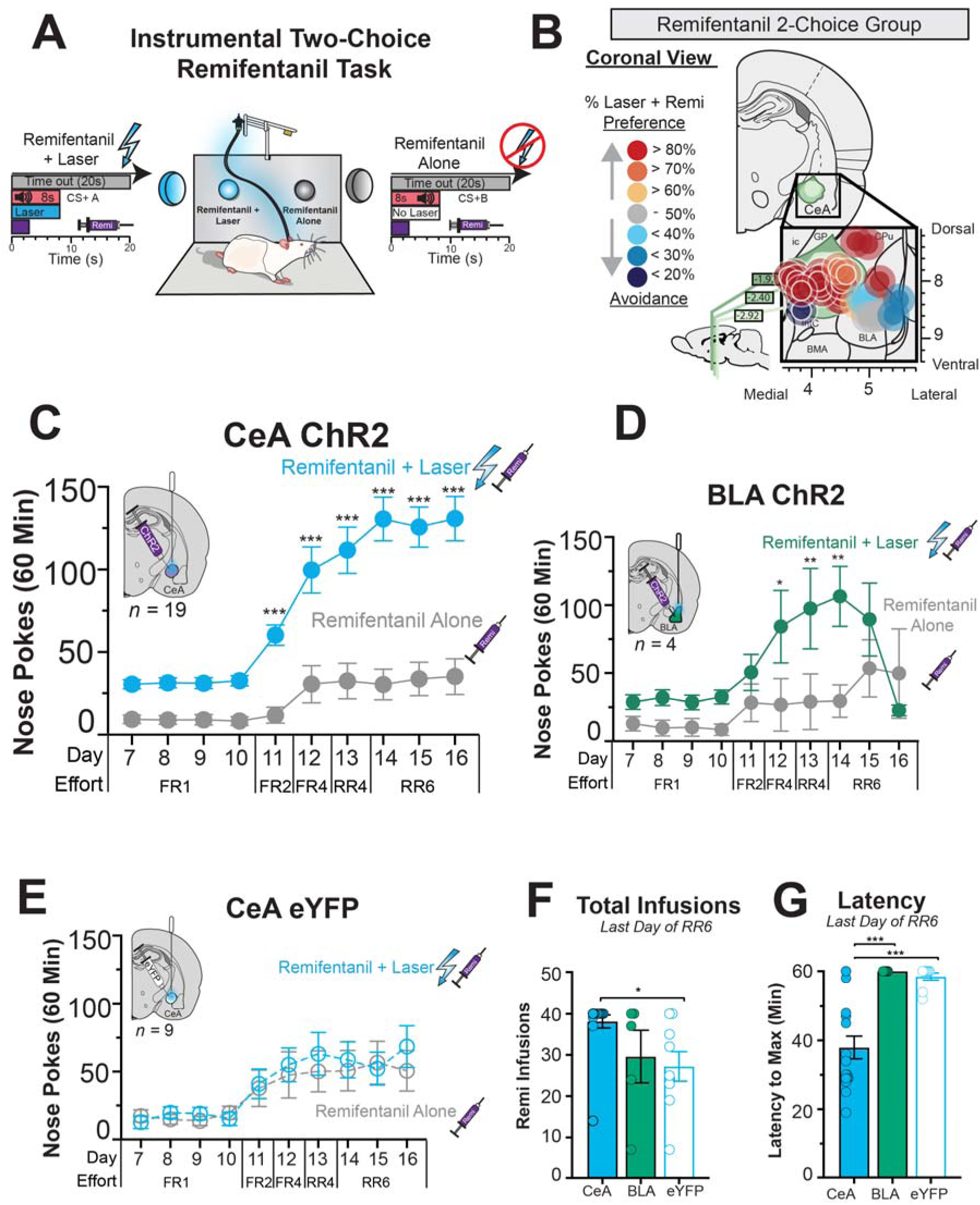
Optogenetic CeA neuron excitations focus motivation for intravenous remifentanil. **A)** Shows instrumental 2-choice task where rats nose poke into two different portholes. Responses to one port earn an intravenous remifentanil infusion (Remifentanil Alone: 2µg/kg; 2.8s duration; FR1-RR6 Schedule) accompanied by a discrete 8s CS+ tone. Nose poking into a second port located on the opposite side of the same wall earned an identical intravenous remifentanil infusion (Remifentanil + Laser: 2µg/kg; 2.8s duration; FR1-RR6 Schedule) paired with a different 8s CS+ tone and additional blue laser stimulation (25 Hz, 1-3 mW, 8s). **B)** Mapped sites of optic fiber implants for each ChR2 rat in remifentanil vs. remifentanil group. Symbol colors in both groups represent the percent preference for an individual rat’s preference for that rat’s laser-paired reward over the non-laser paired reward calculated as the average of the last two sessions. Ic, internal capsule; GP, globus pallidus; CPu, caudate putamen; BLA basolateral amygdala, CeA, central amygdala; IntC, intercalated amygdala; BMA, basomedial amygdala. **C)** ChR2 CeA neuron stimulation captures choice for intravenous remifentanil. CeA ChR2 rats (*n* = 19) singly pursue the laser-paired remifentanil reward (solid blue symbols) and leave the remifentanil alone option (solid grey symbols) relatively ignored. **C)** Basolateral amygdala ChR2 rats (*n* = 4) also show laser-paired remifentanil preference (solid green symbols), although this preference disappeared on the last day of testing. **D)** By contrast, control inactive virus rats (*n* = 9) lacking the ChR2 gene respond equally between remifentanil alone (dotted grey line and symbols) and remifentanil + laser (dotted blue line and symbols). **E)** By the end of 2-choice testing, CeA ChR2 rats self-administer more total remifentanil relative to eYFP controls. **F)** CeA ChR2 also consume their maximum much quicker than BLA ChR2 rats and eYFP controls, terminating their 60-min sessions ∼20 min early. Data shown as mean ± SEM. * *p* < 0.05, ** *p* < 0.01, *** *p* < 0.001, **** *p* < 0.0001.

Both female and male CeA ChR2 rats developed *Laser + Remifentanil* preferences of similar magnitude with no detectable sex differences, on both FR1 schedule days (**Fig S2b**.; Day10; female ratio: 4:1; male ratio: 4.5:1; laser x sex x day interaction: *F*_9,153_ = 1.18, *p* = 0.31, *n =* 9 females, *n =* 10 males) and higher effort days (**Fig. S2c**; RR6 female preference: 3:1; RR6 male preference: 3:1; *F*_9,130_ = 1.33, *p* = 0.22, *n =* 9 females, *n* = 9 males).

*BLA ChR2 group.* 4 of 7 BLA ChR2 rats also showed an initial preference for their *Laser + Remifentanil* option over *Remifentanil Alone*, whereas 2 of 7 avoided *Laser + Remifentanil*, and 1 rat chose equally between it and *Remifentanil Alone* (**Fig S2d**; **Fig 4d.,**FR1: *F*_9,54_ = 2.10, *p* = 0.045, *n =* 7, RR6: *F*_9,27_ = 2.97, *p* = 0.014, *n =* 4). BLA ChR2 individuals that initially chose Laser + Remifentanil had anatomical placements sites of ChR2 virus infections in BLA, with some viral and fiber spread into nearby endopiriform nucleus and piriform cortex. However, for BLA ChR2 rats as a group, the *Laser + Remifentanil* preference disappeared by final day of RR6 testing (*t*_27_ = 1.54, *p* = 0.99. Consequently, the BLA ChR2 group differed from the CeA ChR2 group in preference between the two options (*F*_9,166_ = 4.05, *p* < 0.0001).Overall, BLA ChR2 females consumed more i.v remifentanil overall than males across both options (**Fig S2e**; session day x sex interaction: *F*_9,45_ = 4.17, *p* = 0.006, *n =* 4 females; *n =* 3 males). However, no significant sex difference was detectable in the effects of laser-pairing (laser x session day x brain region interaction*; F*_9,45_ = 1.54, *p* = 0.16; Day 10; female ratio: 5:1; male ratio: 3:1).

By contrast, control CeA eYFP rats with optically-inactive virus chose equally between Laser + Remifentanil and Remifentanil Alone options (**Fig S2f**., **Fig 4e**.; FR1; *F*_9,99_ = 1.22, *p* = 0.29, *n =* 12; 1:1 FR2-RR6: *F*_9,72_ = 0.42, *p* = 0.92, *n =* 9), without showing detectable preference or avoidance of CeA laser (1:1 FR1 (Day 10) Preference Ratio; 1:1 RR6 (Day 16) Preference Ratio). Therefore, the control eYFP group differed from the CeA ChR2 group on both FR1 schedule (laser x session day x virus: *F*_9,261_ = 4.79, *p* < 0.0001) and higher RR6 schedules (*F*_9,211_ = 5.15, *p* < 0.0001).

#### Total opioid intake

CeA ChR2 rats always took the maximum 40 infusions per session, reaching the 80 µg/kg remifentanil maximum dose each day (**Fig 4f**; CeA ChR2: 38.1 ± 1.6; CeA eYFP: 27.2 ± 3.6; BLA ChR2: 29.6 ± 6.4; *F*_2,27_ = 4.42, *p*= 0.02). CeA ChR2 rats overall consumed more total remifentanil than eYFP controls (*t*_27_ = 2.79, *p* = 0.02, *d =* 3.18, CI = 1.8, 20.0). Further, CeA ChR2 rats also reached their maximum 40 infusion ceiling 20 minutes earlier than either eYFP control rats or BLA ChR2 rats (for those in the latter two groups that reached maximum limit) (**Fig 4g**; RR6 Day 16 Session Times; CeA ChR2: 37.9± 3.3; CeA eYFP: 54.4± 1.0; BLA ChR2: 60.0± 0.0; *F*_2,27_ = 16.6, *p* < 0.0001; CeA ChR2 vs eYFP comparison: *t*_27_ = 4.92, *p* = 0.0001, *d =* −1.90, CI = − 30.15, - 8.65; CeA ChR2 vs BLA comparison: *t*_27_ = 4.30, *p* = 0.0006, *d =* −2.36, CI = −34.4, −10.93).

### CeA ChR2 stimulation magnifies the intensity of motivation to seek opioid: Breakpoint task

In a progressive ratio test of incentive motivation intensity, laser excitation increased the effort breakpoint of only CeA ChR2, but not BLA ChR2 or eYFP rats to self-administer i.v. remifentanil, reaching breakpoints in their Laser + Remifentanil conditions (50.0 ± 5.3) that were >190% on average of that in their *Remifentanil Alone* condition (**Fig. 5a**; 26.4 ± 3.7; *F*_2,23_ = 4.48, *p*= 0.02; *t*_23_ = 5.40, *p*= 0.001, *n* = 14). Overall, CeA ChR2 made twice as many nosepokes in the *Laser + Rem*ifentanil condition (201.3 ± 26.2 nosepokes) than in the *Remifentanil Alone* condition (**Fig. 5b**; 92.5 ± 15.5; *F*_2,23_ = 4.61, *p*= 0.02, *t*_23_ = 5.45, *p*= 0.001, *n* = 14).

**Figure 5.**
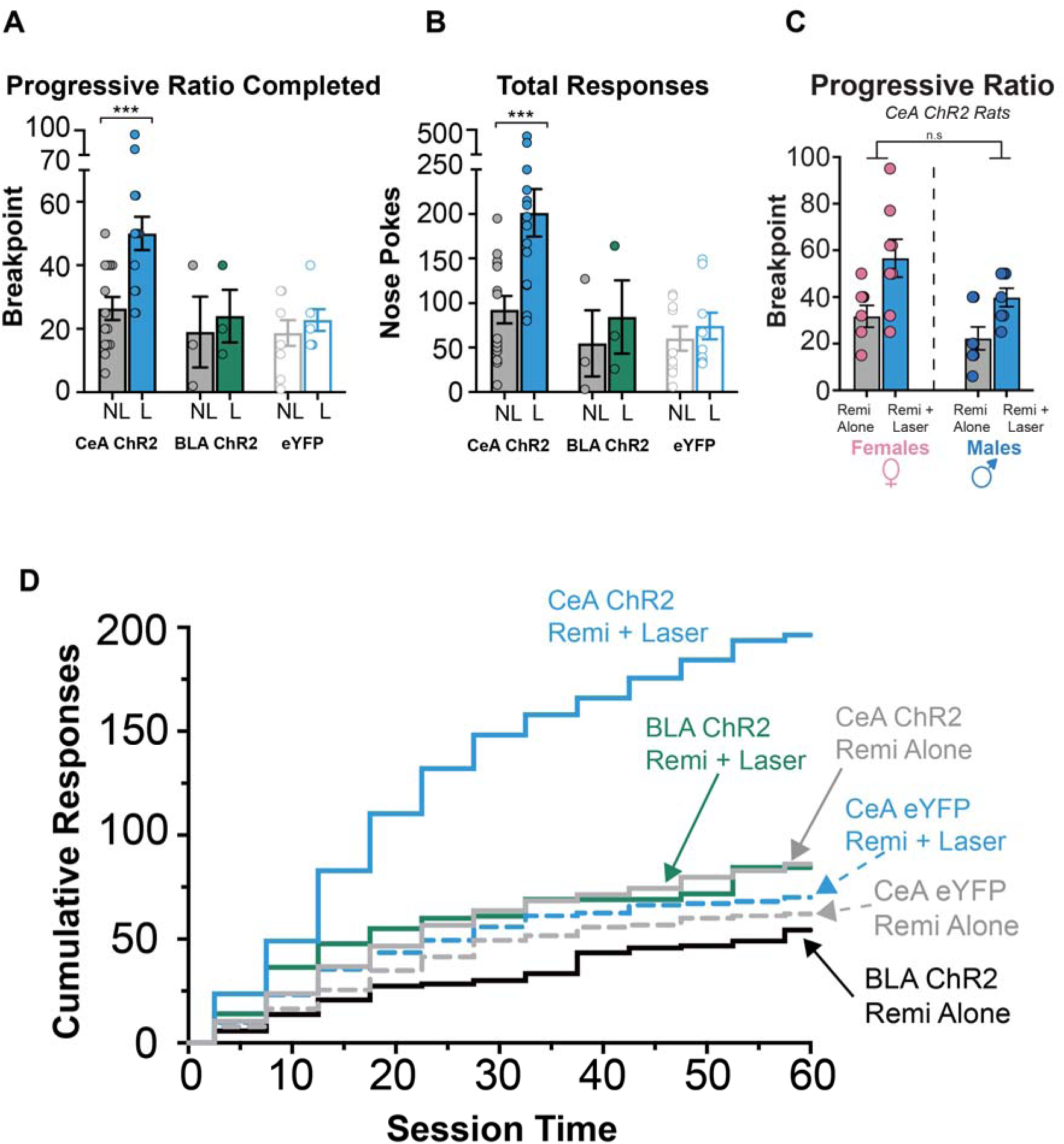
CeA neuron stimulation amplifies breakpoint motivation. We used a progressive ratio test of motivation on two consecutive days. On one day, rats responded for Remifentanil + Laser and on the other day, rats responded for Remifentanil Alone on a progressive ratio schedule of reinforcement so that effort required to obtain the next remifentanil infusion increased exponentially. A) CeA ChR2 rats (*n* = 14) reached higher breakpoints for Remifentanil + Laser and B) made more total responses. By contrast BLA ChR2 rats (*n* = 3) and eYFP controls (*n* = 9) worked equally for remifentanil regardless of laser condition. C) On average, male (*n* = 7) and female (*n* = 8) CeA ChR2 rats worked ∼160% greater for laser-paired remifentanil vs. remifentanil alone during progressive ratio tests and made equal laser-paired remifentanil responses. D) Shows cumulative responses on progressive ratio tests made be CeA ChR2, BLA ChR2 and eYFP control rats. Data are shown as mean ± SEM. *** *p* < 0.001.

This difference became significant within the first 10 min of the sessions, when CeA ChR2 rats had already made 300% more responses in the *Laser + Remifentanil* conditions than in the *Remifentanil Alone* condition (*F*_11,154_ = 22.3, *p*= 0.001, *t*_154_ = 3.61, *p*= 0.005, *n* = 14). Male and female CeA ChR2 rats showed similar laser-induced percentage breakpoint elevations, with no detectable sex difference in magnitude of increase (**Fig. 5c**; Male: Laser + Remifentanil: 39.9 ± 3.9, Remifentanil Alone: 22.3 ± 4.9; Female: Laser + Remifentanil: 56.6 ± 8.1, Remifentanil Alone: 31.8 ± 4.7, *F*_1,13_ = 0.48 *p* = 0.50).

By contrast, control CeA eYFP rats reached similar breakpoints in their *Laser + Remifentanil* condition and *Remifentanil Alone* condition (**Fig. 5a**; NL: 19.9 ± 3.5; L: 22.8± 3.4; *t*_23_ = 0.76, *p*= 0.99, *n* = 9), and made similar numbers of nose pokes on both days (**Fig. 5b**; NL: 60.0 ± 13.7; L: 74.2± 14.9; *t*_23_ = 0.57, *p*= 0.99). BLA ChR2 rats failed to increase breakpoint on the *Laser + Remifentanil* day, and remained similar in effort to their *Remifentanil Alone* day (**Fig. 5a**; NL: 16.5 ± 8.3; L: 26.0± 6.2; *t*_23_ = 0.53, *p*= 0.99, *n* = 3) or in the total number of nose pokes made (**Fig. 5b**; NL: 45.8 ± 27.8; L: 87.3± 29.3; *t*_23_ = 0.69, *p*= 0.99)

## Discussion

Opioid drugs produce distinct euphorigenic and withdrawal effects, and suggestions have arisen that opioid reward and addiction are relatively independent of dopamine-related mesocorticolimbic systems (1, 4–7, 9–11, 56). By such views, motivation for opioid drugs might be expected to resist control by CeA recruitment of mesocorticolimbic circuitry (e.g. activation of ventral tegmentum, substantia nigra, nucleus accumbens, etc.). However, our results show that selective pairing of CeA ChR2 stimulation controlled motivation for opiod vs. sucrose rewards, to direct and amplify incentive salience similarly to studies of other rewards (25). Paired CeA ChR2 stimulation in this study: 1) controlled the target of incentive motivation: determining whether rats pursued sucrose pellets or i.v. remifentanil, depending on which was paired with CeA laser, 2) amplified the intensity of incentive motivation: increasing total opioid intake in the 2-choice task, nearly doubling the intensity of effort breakpoint in the progressive ratio tasks, and tripling the total work amount for laser-paired remifentanil in that task, and 3) narrowed the focus of intensified ‘wanting’: for instance, while CeA ChR2 rats intensely pursued *Laser + Sucrose* they nearly ignored their alternative *Remifentanil Alone.* Conversely, other CeA ChR2 rats specifically pursued *Laser + Remifentanil* and ignored *Sucrose Alone*. Such CeA ChR2-induced control of incentive motivation was reported previously for laser-paired cocaine, alcohol, or sucrose rewards (24, 25, 27, 52), and our results extend this CeA ChR2-induced mesocorticolimbic control to motivation for opioid drugs.

By contrast, control eYFP rats as a group chose roughly equal lower levels between remifentanil and sucrose rewards. In particular, the control eYFP group that had laser paired with remifentanil showed if anything, a marginal preference for their unpaired sucrose (possibly because opioid administration can increase ‘wanting’ and ‘liking’ for palatable foods (57–60). Broader consumption of both options at only moderate levels by control rats is consistent with reports that normal rats often choose sucrose, saccharin, or social alternative rewards over an opioid reward (13–16).

We hypothesize that addictive motivation is more intense and more narrowly focused than ordinary motivation for opioid rewards in non-addicted individuals. Such intensified and focused motivation was induced here by CeA ChR2 recruitment of elevated activation of mesocorticolimbic systems of incentive salience. This mesocorticolimbic motivational amplification and narrowing of focus is consistent with incentive sensitization explanations of how motivation for drugs becomes amplified and focused into addiction via neural sensitization of mesolimbic systems in vulnerable individuals (23, 61, 62). Long-term human heroin users show limbic hyper-reactivity to opioid drug cues in fMRI studies, which can correlate positively with both cue-evoked craving and eventual probability of relapse (63–66). Of course, no addicted human has ever received optogenetic brain stimulations. But the hypothesis that connects our results to incentive sensitization in human addiction is that the final mesocorticolimbic and motivational outcome may be the same, evne though the routes to that outcome are different. That is, CeA ChR2 stimulation recruits a relatively strong mesocorticolimbic activation to intensity and focus ‘wanting’ within minutes to days, whereas a series of smaller endogenous amygdala-related excitations together with sensitization of mesocorticolimbic circuitry in vulnerable individuals may gradually over months to years produce a similar amplified and focused incentive salience outcome in addicted humans (22, 23, 61, 62, 67–69).

*Amplified ‘wanting’, but not ‘liking’*. The CeA ChR2 stimulation that made its paired target more ‘wanted’ did not appear to also make its target more ‘liked’. This issue was approached here when sucrose was the paired target for *Laser + Sucrose* CeA ChR2 rats by applying the affective taste reactivity test (70, 71). This test, which measures homologous orofacial hedonic expressed elicited by sweetness, can detect increases in sucrose ‘liking’ generated by brain manipulations that increase hedonic impact, such as by opioid or optogenetic stimulation limbic ‘hedonic hotspots’ (59, 72–79). The CeA has connections with gustatory sites, such as the brainstem nucleus of the solitary tract (NTS) and parabrachial nucleus of the pons (PBN), or the cortical insula (80–82), and so could conceivably enhance sweetness ‘liking’. However, our taste reactivity results indicate that CeA ChR2 stimulation completely failed to enhance sucrose ‘liking’ reactions in *Laser + Sucrose rats*, despite making the same CeA ChR2 rats intensely ‘want’ laser-paired sucrose. By extension, the selective amplification of opioid pursuit may also have reflected amplified ‘wanting’ but not ‘liking’ for remifentanil in Laser + Remifentanil rats, given that similar mesocorticolimbic Fos elevations were recruited both *Laser + Sucrose* and *Laser + Remifentanil* groups. In short, we conclude that CeA ChR2 pairing made its target more ‘wanted’ here, but not more ‘liked’.

*Not fully explained by laser self-stimulation*. Importantly, laser CeA ChR2 self-stimulation by itself was less potent and reliable in its own as an incentive than its ability to amplify the incentive motivational value of laser-paired remifentanil or sucrose. When assessed at the group level in CeA ChR2 rats, short 1-s laser bin self-stimulation does often occur at statistically significant levels (25, 26, 45, 46, 51–53, 55, 83), though not in all studies (24, 27).

However, self-stimulation is less reliable and often missing when longer 8-s laser bins are used. Here, CeA ChR2 rats initially failed to self-stimulate 8-s laser bins, until after 8-s laser had been associatively paired with either remifentanil or sucrose. That suggests that association with remifentanil or sucrose was required in order for CeA laser to acquire incentive value, rather than intrinsic CeA laser value being responsible for making laser-paired sucrose or remifentanil more ‘wanted’. Further support for that comes from the observation that even CeA ChR2 rats that initially failed to self-stimulate laser at all went on to show as intense selective pursuit of their laser-paired sucrose or laser-paired cocaine in subsequent 2-choice tests as other CeA ChR2 rats that did originally self-stimulate at high rates.

In short, pairing CeA ChR2 stimulation with sucrose, remifentanil, cocaine, etc., appears to produce a synergistic amplification of incentive salience for the targeted unconditional stimulus (UCS), whether or not CeA ChR2 excitation also supports laser self-stimulation in the same rat. The synergy explanation may involve the status of sucrose and remifentanil as hedonically-laden UCSs that themselves activate limbic circuitry (as are cocaine, alcohol, and shock rod). Our results support the hypothesis that the combination of UCS-elicited limbic activation simultaneously with CeA ChR2 stimulation may produce a synergistic recruitment of mesocorticolimbic ‘wanting’ circuitry to generate maximal incentive salience attribution to the paired target (25, 38).

### Conclusion

Opioid addiction is sometimes suggested to differ in mechanisms from other drug addictions, and to rely less on incentive salience mediated by mesocorticolimbic circuitry(1, 4–7, 84). However, our results show indicate that exaggerated and narrowly-focused incentive salience can be controlled for opioid drugs similarly to other rewards, by paired CeA ChR2 stimulations that recruit activation of mesocorticolimbic circuitry (25, 27). Exaggerated incentive salience is not an inherent property of remifentanil, as control eYFP rats sought both remifentanil and sucrose rewards at lower and roughly similar levels, consistent with other reports that ordinary rats often choose alternative sweet or social rewards over opioid self-administration (13–16).

Not all opioid users become addicted, but those who do may have more intense and narrowly-focused incentive motivation to take drugs, similar to the pattern induced here by CeA ChR2 stimulation (12, 15, 20, 21, 23, 62). Addiction is in an individual’s amplified brain and psychological response with other types of addiction, such as excessive ‘wanting’ posited by incentive sensitization.

## Materials and Methods

### Animals

Male and female Sprague Dawley Rats (*n* = 69 total; *n* = 30 males; *n* = 39 females), weighing 250-400 g at surgery were housed in separate-sex rooms at 21°C constant temperature on a reverse 12-h light/dark cycle. All rats had *ad-libitum* access to both food and water through the duration of the experiments. All experimental procedures were approved by the University Committee on the Use and Care of Animal at the University of Michigan.

### Surgery

#### Optogenetic Virus Infusion and Fiber Implant

Rats were anesthetized with isoflurane gas (4-5% at induction; 1-2% maintenance) and received atropine (0.04 mg/kg IP; Henry Schein) before surgery. During surgery, each rat also received subcutaneous injections of cefazolin (100 mg/kg Henry Schein) to prevent infection and carprofen (5 mg/kg; Henry Schein) for pain relief. Cefazolin and carprofen were re-administered 24-h and 48-h post operation. Rats were placed into a stereotaxic apparatus (David Kopf Instruments) and received bilateral 0.75 uL infusions into CeA or BLA of either an AAV ChR2 (AAV5-Hsyn-ChR2-eYFP; CeA *n =* 29; BLA *n* = 7 UNC Vector Core, Chapel Hill, NC) or inactive control virus lacking the ChR2 gene (AAV5-Hsyn-eYFP; *n =* 31; UNC Vector Core, Chapel Hill, NC). Within an individual, CeA placements were bilaterally identical, but coordinates were slightly varied from individual to individual so that sites for the group spanned the entire CeA, ranging from 1.56 – 2.5 (AP), 4.00 – 4.50 (ML), and 7.60-8.00 (DV) all from bregma. The 0.75 µL volume was infused per side over a period of 7.5 minutes, at a constant rate of 0.1 µL per min, and the injector was left in place for an additional 10 minutes to allow for viral diffusion. For optogenetic stimulation, bilateral optic fibers (200 µm) were also implanted 0.3 mm dorsal to the virus infusion site for each rat.

#### Intravenous Jugular Catheter Implantation

A chronic intravenous jugular catheter for intravenous delivery of remifentanil solutions was implanted during a second surgery 3 weeks after the CeA procedure (25, 27, 70, 71, 85). Rats were anesthetized using isoflurane gas similarly to the method described above and administered perioperative treatment for infection, pain, and inflammation. The intravenous (i.v.) jugular catheter was Silastic tubing (id: 0.28 mm; od: 0.61 mm; Plastics1) threaded into the right jugular vein of each rat and secured proximally and distally using non-absorbable polyamide thread (Braum). The outer end of the catheter was passed under the skin of the shoulder, along the dorsal back, and exited from a secured subcutaneous anchor near the midscapular region. Catheter patency was maintained during the initial 10 day post-operative period via daily flushes with 0.2 mL of saline solutions containing 5 mg/mL gentamicin sulfate (Henry Schein) to prevent infection, followed by 0.1 mL of 3.5 mg/mL heparinized saline (Sigma-Aldrich). Subsequently, catheters were flushed daily with either 0.1 mL heparinized saline or 0.2 mL of sterile saline on alternating days. Catheter patency was assessed via intravenous infusion of 0.2 mL methohexital sodium (20 mg/mL; JHP Pharmaceuticals) prior to the start of behavioral testing, roughly midway during the series of behavioral test days, and again at the conclusion of behavioral experiments. Catheters were considered patent if rats became ataxic within 10s of the intravenous injection, and only data from rats with patent catheters were included in the behavioral analysis.

#### Oral Cannula and Optic Fiber Implantation

Given that our initial results indicated that CeA ChR2 pairing controlled ‘wanting’ for sucrose versus remifentanil targets, it was important to ask whether CeA ChR2 stimulation also enhanced hedonic impact or ‘liking’ of a paired stimulus. This question could be answered at least for paired sucrose via the affective taste reactivity test, based on sweetness-elicited orofacial expressions of ‘liking’ (e.g. lateral tongue protrusions) versus ‘disgust (e.g., gapes) (70, 71, 85, 86). A separate group of rats first received an AAV5 ChR2 virus in CeA as above (AAV5-hsyn-ChR2-eYFP, *n =* 6 or AAV5-hsyn-eYFP control, *n =*6, UNC Vector Core), except that optic fibers were not implanted during the initial surgery. Following infusion of the optogenetic virus, the head wound was sutured, and rats were allowed to recover for at least three weeks and to allow time for virus expression. A second surgery was then performed to implant oral cannula and CeA optic fiber. Rats were re-anesthetized with isoflurane as described above for implantation of intracranial optic fibers and of bilateral oral cannulas, which allowed for direct oral infusions of sucrose, and quinine solutions. Each oral cannula (polyethylene-100 tubing) entered the upper cheek just lateral to the secondary maxillary molar, ascended beneath the zygomatic arch, and exited the skin at the dorsal head, where it was secured with skull screws and a dental acrylic headcap. In the same surgery, rats were implanted with bilateral optic fibers (200 µm), aimed to place each fiber tip 0.3 mm dorsal to the rat’s bilateral virus microinjection sites, and anchored with the same acrylic headcap. Cefazolin and carprofen were again administered and repeated post-operatively as above. All rats were allowed to recover for 1 week prior to behavioral testing.

### Behavioral Testing

#### Experiment 1: Remifentanil vs. Sucrose 2-Choice Task

CeA ChR2 rats (*n* = 10; *n* = 6 females, *n* = 4 males; CeA eYFP: *n =* 13*; n =* 9 females, *n =* 4 males) were used to assess whether pairing CeA ChR2 excitation could control choices between intravenous remifentanil versus natural sucrose pellets. Laser parameters were based on previous published studies with cocaine and our remifentanil own pilot tests (Baumgartner et al., 2022; Warlow et al., 2017). Half the group of CeA ChR2 rats were arbitrarily and permanently assigned to choose between *Laser + Sucrose* versus *Remifentanil Alone* (2 µg/kg), by making nosepokes into either of two portholes, each porthole permanently assigned to earn its designated reward (*n* = 3 females, *n* = 2 males). Sucrose pellets were delivered into a recessed dish located in the chamber wall between the two ports. Remifentanil i.v. infusions were delivered by a syringe pump attached to the rat’s i.v. catheter (2 µg/kg in 50 µL of 0.9% isotonic saline; delivered over 2.8-s). This *Laser + Sucrose* group had 8-sec laser selectively paired with choosing sucrose when they made nosepokes into the Laser + Sucrose porthole. By contrast, for these rats, remifentanil was earned alone without laser when they made nosepokes into the remifentanil porthole. Each rat received the same assignment throughout the experiment. Once a reward was delivered, both nose ports retracted into the back wall for a 20-s time out intertrial period. During this time out, no responses could be made, and no reward was available.

The other CeA ChR2 group (*n* = 3 females, *n* = 2 males) was assigned to choose between *Laser + Remifentanil* versus *Sucrose Alone,* again by making nosepokes into either of two permanently assigned portholes. For all CeA ChR2 rats, nose pokes into the porthole that earned laser-paired reward delivered photoexcitation (1-3 mW, 25 Hz, 8-s pulse train) that began with the final nosepoke and continued to overlap with receipt of its paired sucrose or remifentanil reward.

Instrumental training and choice sessions were conducted in MedAssociates operant chambers with clear Plexiglas floors and two retractable portholes. Infusion pumps were located outside each chamber to allow for intravenous delivery of remifentanil. In initial training sessions (prior to making choices), rats were initially trained for several days with a single outcomes in a 60-min session, then alternating to their other outcomes on the next day. Training continued alternating each outcome daily until each rat had cumulatively earned a total of >50 sucrose pellets and >50 remifentanil infusions (either with laser or without depending on their group assignment). This was done to thoroughly acquaint rats with the relative value of each of their outcomes, to help guide subsequent choice. During single-outcome training sessions, only one of the two portholes or one of the two levers was inserted into the box per day. Each manipulandum was permanently assigned to earn its own particular reward type (i.e., *Laser + Sucrose; Laser + Remifentanil; Sucrose Alone or Remifentanil Alone*). On the next day, the other porthole or lever was solely available, and nosepokes in it earned the alternative reward.

*2-Choice tests: Days 1-9*. After initial training was completed, 9 days of 2-choice test sessions were presented. Each 2-choice session began with a single choice trial in which only one porthole was randomly inserted until a nose poke was made and its customary reward earned (either sucrose or remifentanil; accompanied by laser or not depending on individual assignment). The first porthole was then retracted, and a 20-s time out intervened before the alternative porthole was inserted and it remained until the rat earned its alternative reward. This single-choice cycle was repeated once more until each rat had earned two remifentanil rewards and two sucrose rewards on each day, with one reward type accompanied by laser and the other not depending on individual assignment. These initial single-choice presentations were delivered to remind the rat of each of its two outcomes at the beginning of the day, before the rat was allowed to choose freely between the outcomes for the rest of the session. In the remaining 2-choice trials, both portholes or levers were always inserted simultaneously, allowing rats to freely choose between intravenous remifentanil or sucrose. Upon a nosepoke into a chosen porthole, that porthole’s customary reward was delivered accompanied by an assigned auditory cue that identified choice of sucrose or remifentanil (either tone or white noise; assignment counterbalanced across rats). Laser illumination (8-sec) also occurred if the rat had chosen its assigned laser-paired reward. Both portholes were retracted for a 10-min time out, before both portholes were again inserted for the next trial to allow a 2-choice opportunity again. The 2-choice/timeout combination was repeatedly presented again for the remainer of the 2-hr sessions, allowing up to 12 independent choices per sessions. These 2-chocie sessions were repeated daily for 9 days.

### Preference persistence over Laser Extinction: Days 10-14

To assess whether a CeA ChR2-induced focus of motivation would persist after paired CeA neuronal excitation was discontinued, we tested a previously induced preference for sucrose vs remifentanil would persist when laser was discontinued over 5 daily consecutive laser-extinction sessions. In each 2-hr laser-extinction trial, 2-choice presentations were offered similarly to above, but neither reward triggered CeA laser illuminations, and so both sucrose and remifentanil were earned without laser for all rats.

### Laser Self-Stimulation Tests

We assessed the incentive value of ChR2 CeA stimulation on its own, in the absence of either remifentanil or sucrose. Each rat above was also tested for laser self-stimulation in two self-stimulation tasks. Rats were tested for both active-touch and place-based laser self-stimulation before any 2-choice training/test sessions began, and retested again after the conclusion of all 2-choice tests.

Active-touch Self-Stimulation Task. In an active object-touch task, rats could earn laser illuminations in 30 min sessions by instrumentally touching a designated empty metal rod or drinking spout (active rod/spout) that protruded from the backwall of a Med Associates chamber (a different chamber from that used in 2-choice tests). A second rod or spout was placed 10 cm apart and earned nothing (inactive rod/spout), serving simply as a baseline measure of exploratory touches. The floor of the chamber was a metal grid floor, enabling a circuit to detect contacts onto each respective spout or rod. Each touch on the designated laser rod/spout (permanently assigned for each rat; assignments balanced across rats) delivered a brief CeA laser illumination (1-3 mW; 25 Hz) of either 1-sec bin duration or 8-sec duration in different sessions (i.e. identical to laser bin duration in 2-choice task). Each 1-sec and 8-sec duration was tested in 3 consecutive daily sessions, with order of 1-sec vs 8sec tests balanced across rats.

Place-based Self-Stimulation Task. In a separate place-based self-stimulation task, rats could earn laser illuminations (25 Hz; 3-s pulse trains) by entering a designated one of two chambers of a two-chamber Plexiglas apparatus, and earn repeated laser illuminations by simple remaining there (76 cm x 38 cm x 38 cm). Each chamber was distinguished by its own set of visual cues (striped vs dotted walls) and tactile cues (small grid vs. large grid floor). A pre-test was conducted to assess chamber preference, in which rats were allowed to move freely between the two compartments without any laser stimulation. After this pre-test one chamber was randomly assigned (non-biased design) as the laser-delivering chamber (assignment balanced across rats). Rats could earn laser illumination by entering their laser-delivering compartment (1-3 mw; 25 Hz; 3-s pulse trains). Laser illuminations continued to cycle (3-s ON/8-s OFF) as long as the rat remained in that chamber. Exiting that chamber terminated laser delivery, and entry into the alternative no-laser chamber ensured absence of laser stimulation. For each rat, the same chamber was assigned as its laser-delivering chamber for three consecutive daily 15-min sessions.

### Taste Reactivity Testing

CeA ChR2 rats that had earned *Laser + Su*crose (vs *Remifentanil Alone*) in the 2-choice task were subsequently assessed for CeA ChR2 excitation effects on sweetness ‘liking’ in the affective taste reactivity task. Each rat was habituated to the transparent plastic taste reactivity test chamber for 30 minutes on four consecutive days before any behavioral testing occurred. On the last two days of habituation, rats received oral infusions of a 0.03M sucrose solution to habituate them to infusion of fluid into the mouth. In subsequent taste reactivity tests, affective orofacial reactions (i.e., positive ‘liking’ versus negative ‘disgust’ patterns) elicited by oral infusions of sucrose solutions (0.03M), or quinine (3 x 10^-4^ M), were videorecorded for offline analysis (70, 71). Orofacial reactions were videorecorded through a close-up lens facing an angled mirror underneath the transparent floor, positioned to capture a clear view of the mouth and face. Taste solutions (1 ml) were delivered into the mouth of rats through PE-50 tubing connected to a PE-10 delivery nozzle, at a constant 1ml/min rate during the 1 min infusion, via a syringe pump, connected to the oral cannula. Only one type of taste was tested per day (sucrose, water, or quinine) for a total of 2 x 1mL infusions per day, either with laser stimulation, or without laser as a within-subject baseline. During laser trials, laser illumination (1-3 mW; 25 Hz, 15 ms pulse/cycled 5-s ON/5-s OFF) throughout the 1-min test. Each laser parameter was tested at least twice for each rat in separate daily tests.

Taste reactivity videos were scored subsequently for positive hedonic ‘liking’ reactions, aversive ‘disgust’ reactions, and neutral taste reactions in slow-motion speeds ranging from frame-by-frame to 1/5^th^ normal speed using The Observer Software (Noldus; Leesburg, VA). Positive hedonic or ‘liking’ responses were considered to be; lateral tongue protrusions, paw licks, and rhythmic midline tongue protrusions. Aversive ‘disgust’ reactions were: gapes, forelimb flails, headshakes, face washes, chin rubs, and paw treading. Neutral responses (i.e., relatively uncoupled from hedonic impact) were: passive dripping of solution out of the mouth, rhythmic mouth movements, and grooming. A time-bin scoring system was used to ensure each type of affective reaction contributed equally to the overall affective score (70, 73, 74). Rhythmic mouth movements, paw licks, passive dripping, and grooming were all scored in 5-s time bins, because these behaviors typically are emitted in bouts of relatively long duration. Any emissions of these behaviors up to 5-sec was counted as a single occurrence; emissions of 5-sec to 10-sec counted as two occurrences, etc. Midline tongue protrusions and pat-treading were scores similarly, but in 2-s bins, as typically emitted in shorter bouts. Lateral tongue protrusions, gapes, flails, headshakes, and chin rubs were counted as discrete events every time they occurred, because these can occur singly or in several brief repetitions. A total positive hedonic (i.e., ‘liking’) score was then calculated by combining component scores of rhythmic tongue protrusions, paw licks, and lateral tongue protrusions. A total negative aversive (i.e., ‘disgust’) score was calculated by combining gapes, forelimb flails, head shakes, paw treading, face washes, and chin rubs (70, 74, 78).

### Experiment 2: CeA ChR2 amplification of remifentanil incentive value

Previous studies showed that CeA ChR2 stimulation amplified the incentive value of laser-paired cocaine or sucrose in addition to narrowing the focus of ‘wanting’ to that target. Here we assessed in two ways whether CeA ChR2 pairing similarly amplifies the incentive value of laser-paired remifentanil over the value of remifentanil by itself. First, we used a two-choice task similar to the one used previously for i.v. cocaine to assess if CeA ChR2 pairing caused *Laser + Remifentanil* to be persistently sought while ignoring *Remifentanil Alone* (27). Second, we used a progressive ratio test (i.e., breakpoint) task to ask whether CeA ChR2 stimulation would increase the effort rats were willing to expend to earn *Laser + Remifentanil* compared to their efforts for *Remifentanil Alone*.

*Two-Choice Task*. In this task, a new group of CeA ChR2 rats or eYFP control rats could choose between two nosepoke options that delivered either *Laser + Remifentanil* or *Remifentanil Alone* (ChR2: *n =* 8 females, *n =* 10 males; eYFP: *n =* 4 females, *n =* 5 males; 1-2 mW; 25 Hz, 15-ms pulse; 8-s pulse train)*. Remifentanil Alone* or *Laser + Remifentanil* infusions were each accompanied by their own distinctive 8-sec auditory cue to provide an additional signal of which reward was earned (either tone or white noise; assignment counterbalanced across rats). The very first session was 2-h in duration, and all subsequent sessions were 1-h. Besides automated beam-break measures of nose-poke responses, two video cameras, one placed below the transparent floor and the other mounted on a chamber wall recorded behavior for subsequent off-line video analysis.

Each 2-choice session began with a series of 4 single choice presentations, in which only one porthole was presented to earn its customary outcome: either Remifentanil Alone or Laser + Remifentanil. Rats had to successfully poke into the available nose port until intravenous delivery of the drug was earned together with its customary laser or no-laser condition. After a 20-s timeout period, the alternative nose port was presented until the rat earned the alternative reward, either Laser + Remifentanil or Remifentanil Alone. Then the sequence of first porthole followed by second porthole was repeated once more, so that each outcomes earned twice. These single-choice trials were conducted at the beginning of every sessions to ensure that rats experienced both types of remifentanil outcomes each day before having to choose between them.

Both portholes were presented simultaneously for the rest of each session to allow choices between them. After each outcome was chosen and received, the portholes retracted for a 20-sec time out period. Then simultaneous presentation of both portholes was repeated for the next choice. This continues for 1-hr or until the rat earned a maximum ceiling of 40 opioid infusions. Sessions were repeated daily for 1-days on a Fixed Ratio (FR1) schedule, where each nose poke resulted in an intravenous remifentanil infusion. Subsequently the fixed ratio effort requirement was increased over Days 11-16 to assess the robustness of any preference in the following order: FR2, FR4, RR4, RR6, RR6, RR6.

### Is Laser Itself the Goal? Laser Self-stimulation Test

A second way to assess if laser CeA ChR2 stimulation was rewarding by itself and added a separate value was to assess laser self-stimulation by itself in a new task not associated with remifentanil. In novel chambers, the active-response laser self-stimulation task described in Experiment 1 was used. CeA ChR2 rats could earn a brief laser illumination (15 ms laser pulses; 1-3 mW; 25 Hz; either 1-s or 8-s duration bin depending on trial day). Each combination of laser intensity/duration parameters was repeated on 3 consecutive days of self-stimulation (30-min sessions, order of combinations balanced across rats).

### Progressive Ratio Test of Effort Breakpoint: Does CeA ChR2 Stimulation Amplify Intensity of Incentive Motivation for Remifentanil?

To assess if laser increases the magnitude of incentive motivation for remifentanil, the same CeA ChR2 rats subsequently underwent two days of progressive ratio (PR) testing after all 2-choice tests had finished. On one PR day (order balanced across rats) only the *Remifentanil Alone* port was available to earn its customary opioid reward without laser. On the other PR day, only the *Laser + Remifentanil* porthole was available, which earned its customary opioid reward together with 8-sec CeA Chr2 laser stimulation. Within each PR session, the number of nose pokes needed to earn the next remifentanil infusion increased exponentially after each delivery according to a progressive ratio schedule of reinforcement with the following formula [PR= [ 5*e*^(reward^ ^number^ ^x^ ^0.2)^]-5 rounded to the nearest integer] (24, 27). Work requirements for this schedule were 1,2, 4, 6, 9, 12, 15, 20, 25, 32, 40, 50, 62, 77, 95, 118, etc. The effort breakpoint reached when responding stopped was compared between laser vs no-laser conditions (i.e., the maximum effort price paid in terms of responses to earn remifentanil within each 1-hr session).

## Statistical Analysis

Remifentanil 2-choice sessions, sucrose vs. remifentanil, spout self-stimulation, and place-based self-stimulation tasks were analyzed by mixed ANOVA, followed by t-tests with Bonferroni corrections for individual comparisons. Friedman’s two-way ANOVAs were used for non-parametric tests, following Wilcoxon sign-ranked tests. Significance was set at *p*< 0.05

## Acknowledgments

We thank Nina Mostovoi, Madeleine Granillo, and Carina Castellanos for technical assistance, and Marc Bradshaw for equipment construction.

## Author Contributions

IM: Conceptualization, Methodology, Investigation, Supervision, Visualization, Writing

DN: Methodology, Investigation, Visualization, Writing

KB: Conceptualization, Methodology, Investigation, Supervision, Visualization, Writing

---

**Figure S1.**
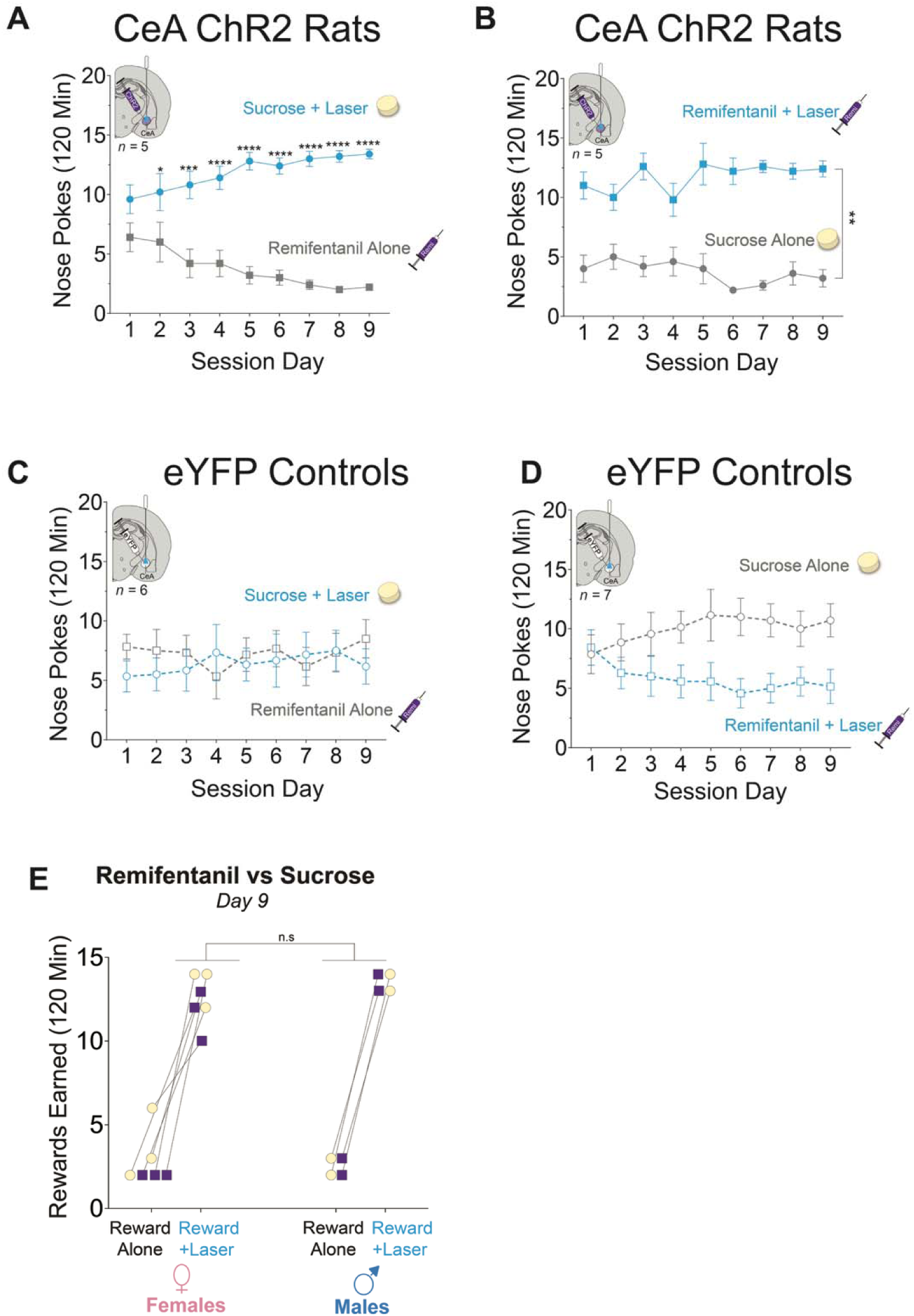
CeA ChR2 laser pairing controls pursuit of remifentanil versus sucrose. **A-D)** Nose pokes by rats in each group during each remifentanil vs. sucrose choice session. **A)** Sucrose + Laser CeA ChR2 rats (*n* = 5) exclusively poke for sucrose + laser and ignore intravenous remifentanil. **B)** Remifentanil + Laser CeA ChR2 rats (*n* = 5) singly pursue intravenous remifentanil and ignore sucrose. **C,D)** Neither group of eYFP controls (Sucrose + laser: *n* = 6; remifentanil + laser: *n* = 7) shows preference for either sucrose or remifentanil. **E)** Rewards earned by individual female (left) and male (right) ChR2 rats during the last remifentanil vs. sucrose choice session. Male (laser preference ratio 5:1; *n* = 4) and female (laser preference ratio 4:1; *n* = 6). Sucrose responses in yellow circles and remifentanil responses in purple squares. All data presented as mean and SEM. * *p* < 0.05, ** *p* < 0.01, *** *p* < 0.001, **** *p* < 0.0001.

**Figure S2.**
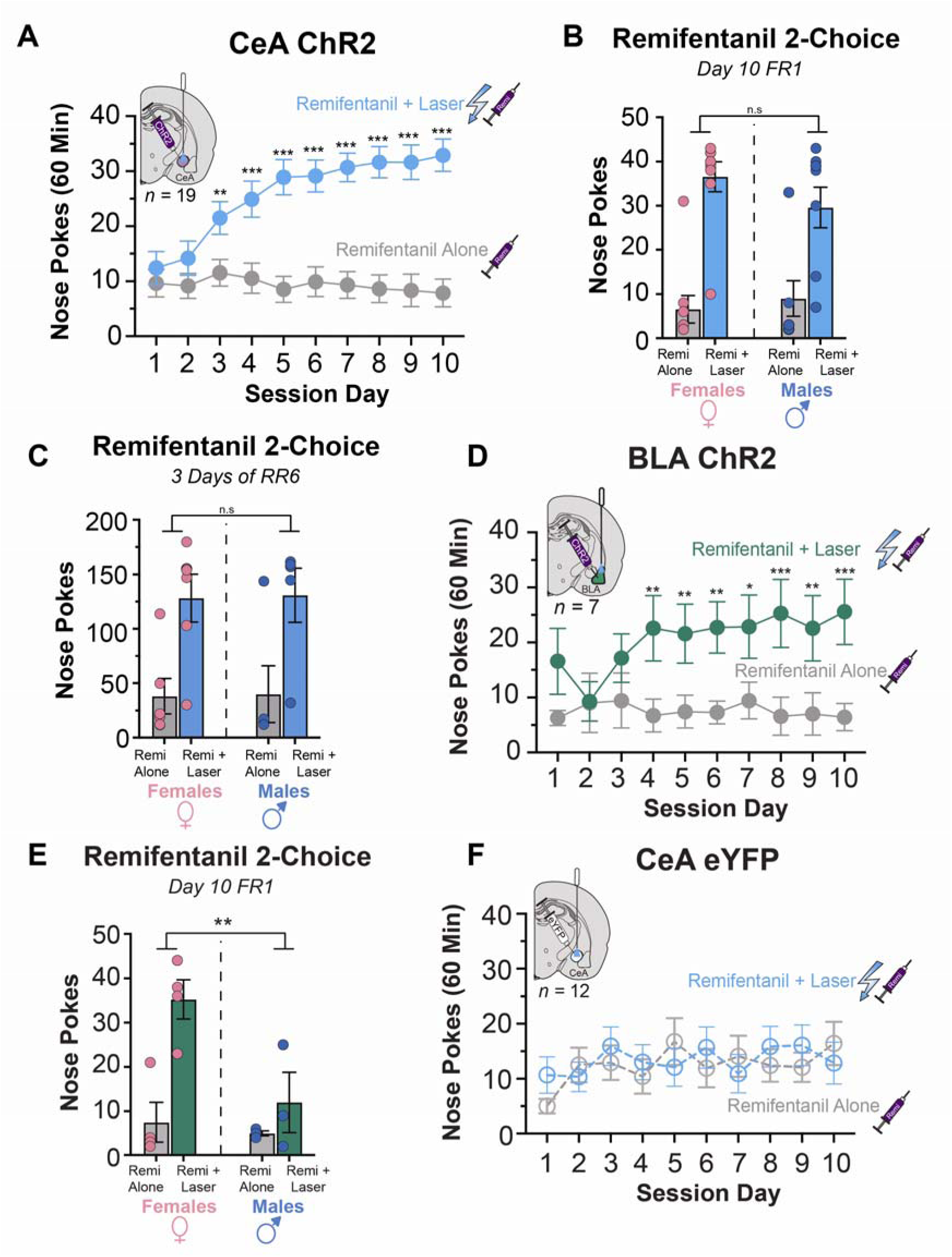
CeA ChR2 neuron stimulation captures choice for intravenous remifentanil. **A)** CeA ChR2 rats (*n* = 19) almost exclusively pursue intravenous remifentanil reward paired with optogenetic CeA neuron excitation (solid blue lines) compared to intravenous remifentanil alone (solid grey lines) during initial 10 days of FR1 testing. **B)** No sex differences detected between male and female CeA ChR2 rats during instrumental choice sessions. Male (*n* = 10) and female (*n* = 9) rats equally preferred laser-paired remifentanil over remifentanil alone during remi vs. remi test sessions. Data is from Day 10 of FR1 choice sessions. **C)** Male (laser preference ratio 3:1; *n* = 5) and female (laser preference ratio 3:1; *n* = 6) CeA ChR2 rats equally prefer laser-paired remifentanil over remifentanil alone during 2-choice test sessions. Data presented by averaging last three RR6 sessions. **D)** BLA ChR2 rats also showed preference for laser-paired remifentanil (solid green lines) compared to remifentanil alone (solid grey lines). **E)** Female BLA ChR2 (*n* = 4) rats earned more total i.v. remifentanil compared to male BLA ChR2 rats (*n* = 3). **F)** eYFP control rats (*n* = 12) chose equally between intravenous remifentanil alone (dotted gray lines) and laser-paired remifentanil (dotted blue lines). Data are shown as mean and SEM. * *p* < 0.05, ** *p* < 0.01, *** *p* < 0.001, **** *p* < 0.0001.

